# Amino acid intake strategies define pluripotent cell states

**DOI:** 10.1101/2022.11.16.516803

**Authors:** Pavlina K. Todorova, Benjamin T. Jackson, Vidur Garg, Katrina I. Paras, Yanyang Chen, Sanjeethan C. Baksh, Jielin Yan, Anna-Katerina Hadjantonakis, Lydia W. S. Finley

## Abstract

Mammalian pre-implantation development is associated with striking metabolic robustness, as embryos can develop in a wide variety of nutrient conditions including even the complete absence of soluble amino acids. Here, we show that mouse embryonic stem cells (ESCs) capture the unique metabolic state of pre-implantation embryos and proliferate in the absence of several essential amino acids. Amino acid independence is enabled by constitutive uptake of exogenous protein through macropinocytosis alongside a robust lysosomal digestive system. Upon transition to more committed states, ESCs reduce digestion of extracellular protein and instead become reliant upon exogenous amino acids. Accordingly, amino acid withdrawal selects for ESCs that mimic the pre-implantation epiblast. More broadly, we find that all lineages of the pre-implantation blastocysts exhibit constitutive macropinocytotic protein uptake and digestion. Together, these results highlight exogenous protein uptake and digestion as an intrinsic feature of pre-implantation development and provide insight into the catabolic strategies that enable embryos to sustain viability prior to implantation.

## Introduction

Mammalian development is initiated by a series of precisely orchestrated cell divisions that first cleave the fertilized zygote into multiple blastomeres that then divide to form the three lineages of the pre-implantation blastocyst. Upon implantation, the blastocyst initiates a period of explosive cell division and lineage differentiation that enables gastrulation^1^. In the pre-implantation period, cell divisions are reductive; only upon implantation does the embryo begin to gain biomass volume^2^. While even reductive divisions require the production of DNA, cell membranes and newly synthesized proteins, such biosynthetic demands necessarily increase when embryos begin to increase in size. Accordingly, the metabolic strategies used to support proliferation are likely to change dramatically upon implantation. Nevertheless, despite the importance of these early cell divisions for organismal development, little is known about the metabolic networks that define and enable these early embryonic states.

Pioneering studies on mammalian pre-implantation embryos have identified a wide range of culture conditions that support robust development from zygote through late blastocyst stage^3-6^. The capacity for pre-implantation development to proceed in a variety of nutritional environments raises the possibility that pre-implantation embryos harbor significant metabolic plasticity that provides robustness by insulating early development from nutritional fluctuations. Strikingly, although soluble amino acids are the major source of biomass for most proliferating cells^7,8^ embryos can develop to the blastocyst stage in the complete absence of soluble amino acids^3-5^. How this amino acid independence is achieved, and whether it defines pre-implantation states, remains unknown.

Pluripotent embryonic stem cells (ESCs) provide a powerful model system for uncovering links between metabolism and development^9,10^. When cultured with serum and leukemia inhibitory factor (LIF, hereafter S/L), ESCs are heterogeneous and metastable, with a portion of cells exhibiting transcriptional profiles reminiscent of the pre-implantation epiblast lineage, which ultimately gives rise to all 3 germ layers^11^. Addition of inhibitors against MEK and GSK3β (‘2i’) captures ESCs more uniformly in the ‘naïve’ or ‘ground state’ of pluripotency^12,13^. Withdrawal of 2i/LIF (2i/L) allows cells to dismantle the naïve pluripotency transcriptional network and acquire differentiation competence, and addition of fibroblast growth factor 2 (FGF2) and Activin A will drive ESCs to a more developmentally advanced state that represents the post-implantation epiblast (EpiLC)^14-16^. Using these media formulations, we and others have identified intracellular metabolic networks that favor maintenance of the naïve pluripotent state^17-21^. Here, we leveraged these tools to identify the nutrient acquisition strategies that define pre-implantation states.

## Results

### Naïve ESCs proliferate in the absence of essential amino acids

Despite the discovery over 50 years ago that mammalian pre-implantation embryos can develop to the blastocyst stage in the complete absence of soluble amino acids, how embryos sustain cell division and development without a continuous supply of essential amino acids remains unknown. To investigate whether amino acid independence is a hallmark of pre-implantation development, we first assessed amino acid requirements using naïve ESCs as a model of the pre-implantation epiblast lineage. We and others previously demonstrated that the ability to tolerate glutamine withdrawal distinguishes naïve ESCs from their more developmentally advanced counterparts^18-20^ (Supplemental Fig. 1A), but as cells can synthesize this non-essential amino acid, the cellular response to glutamine deprivation is likely to be different than the response to essential amino acid deprivation. We therefore withdrew the essential amino acid leucine, which is the most abundant amino acid in both mouse and human proteomes^22^. Consistent with the essential role of leucine in most proliferating cells, metastable, S/L-cultured ESCs rapidly lost viability following leucine withdrawal (Fig. 1A). In contrast, S/L+2i-cultured ESCs representing the naïve, ground state of pluripotency maintained viability and colony morphology even in the absence of leucine (Fig. 1A, B).

**Fig. 1.**
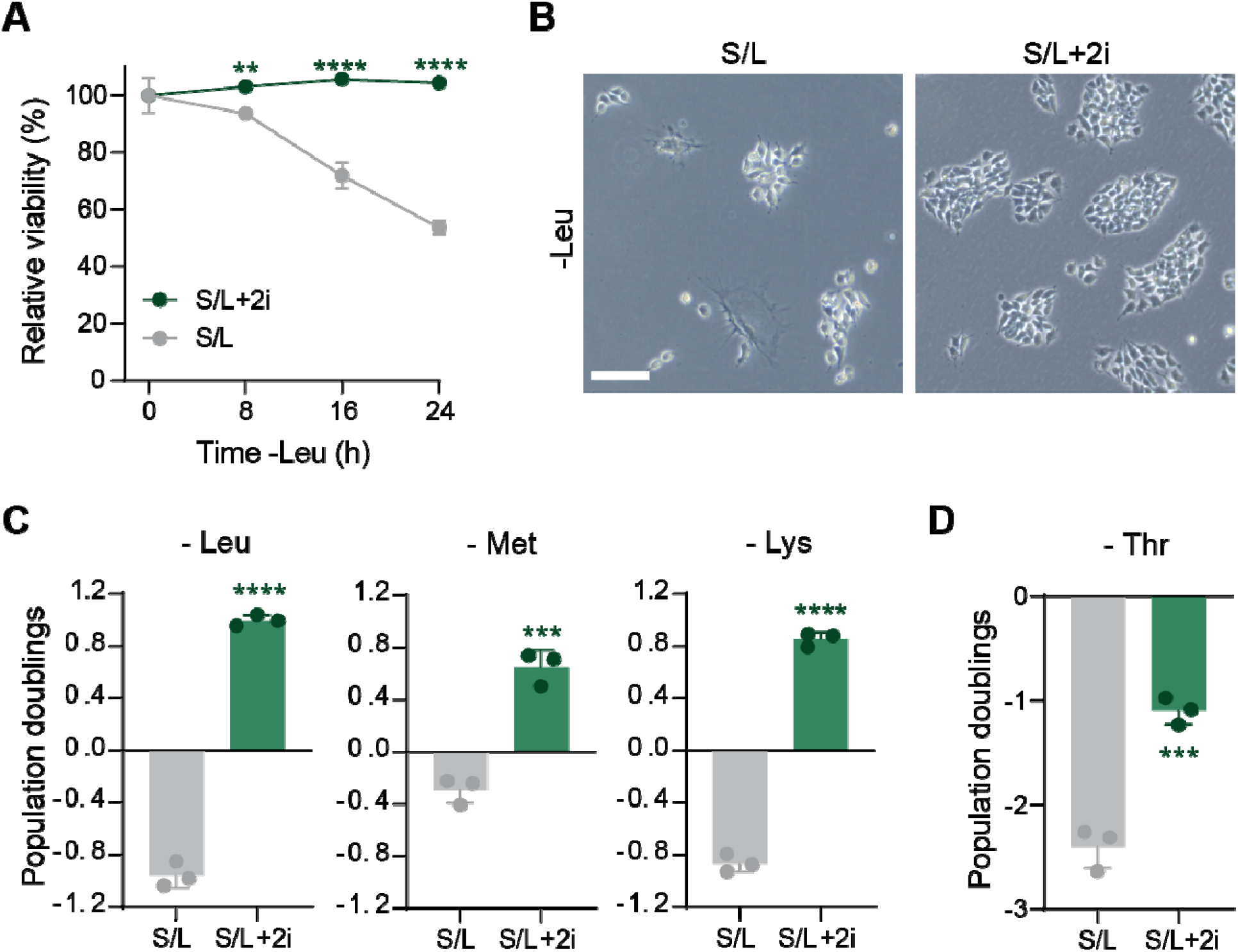
Naïve ESCs can proliferate without essential ammino acids. (**A**) Relative viability (measured by PI exclusion) of S/L and S/L+2i cultured ESCs subjected to leucine deprivation for the indicated times. (**B**) Representative brightfield images of S/L and S/L+2i cultured ESCs after 48 h of leucine withdrawal. Scale bar, 100 µm. (**C**) Population doublings of S/L and S/L+2i cultured ESCs subjected to leucine (Leu), methionine (Met) or lysine (Lys) deficient conditions for 48 h. (**D**) Population doublings of S/L and S/L+2i cultured ESCs subjected to threonine (Thr) deficient conditions for 48 h. Data are mean ± SD, n=3 independent samples. Significance was assessed by two-way ANOVA with Sidak’s multiple comparisons post-test [(A)] and unpaired two-tailed *t* test [(C), (D)] (**p<0.01, \*\*\**p*<0.001, \*\*\*\**p*<0.0001).

To test whether leucine is required for cell division, we quantified changes in cell number following leucine withdrawal. Despite the absence of leucine, two independently derived ESC lines cultured in S/L+2i increased overall cell number, whereas their S/L-cultured counterparts declined in cell number when deprived of leucine (Fig. 1C and Supplemental Fig. 1B). This phenotype was not unique to leucine: S/L+2i-cultured ESCs, but not S/L-cultured ESCs, also proliferated in the absence of methionine and lysine (Fig. 1C). In contrast, both S/L- and S/L+2i-cultured ESCs were dependent on exogenous threonine (Fig. 1D). These results are consistent with previous reports that mouse pluripotent stem cells have unusually high threonine catabolism driven by cell-type specific expression of threonine dehydratase that renders ESCs dependent upon a supply of extracellular threonine^23,24^. Together, these results indicate that naïve ESCs, like pre-implantation blastocysts, can tolerate withdrawal of even essential amino acids so long as the consumption of the amino acid does not outstrip supply.

We next asked whether naïve ESCs tolerate amino acid withdrawal because they can engage in reductive cell divisions (i.e., lacking net gain in volume), which could enable increased cell number without a concomitant gain in biomass volume. Indeed, cell volume decreased following leucine withdrawal in both S/L and S/L+2i-cultured ESCs (Supplemental Fig. 1C). Nevertheless, this decrease in cell volume was more than offset by the increase in cell number when cells were grown with 2i; accordingly, S/L+2i-cultured ESCs were able to increase total biomass volume despite the absence of an essential amino acid (Supplemental Fig. 1D). Because biomass generation depends on continuous availability of all proteinogenic amino acids, these results demonstrate that naïve ESCs have nutrient uptake strategies beyond the conventional uptake of soluble amino acids from the culture medium.

### Naïve ESCs constitutively engage macropinocytosis

Macropinocytosis has emerged as a mechanism for select cancer cells to derive amino acids from extracellular proteins when soluble amino acids are limiting^25-28^. Following nonselective endocytosis, extracellular proteins are delivered to the lysosome where they are degraded into amino acids that can then fuel intracellular metabolic networks^8,29^. This process can be monitored using a form of bovine serum albumin conjugated to a self-quenched fluorescent dye (DQ-BSA) that becomes unquenched following lysosomal hydrolysis^30^. DQ-BSA fluorescence was readily visible in both S/L and S/L+2i-cultured ESCs, and the number of puncta was significantly increased when cells were cultured in the presence of 2i (Fig. 2A, B). Flow-cytometry-based quantification confirmed that 2i-cultured ESCs harbor significantly increased DQ-BSA signal relative to their control counterparts (Fig. 2C).

**Fig. 2.**
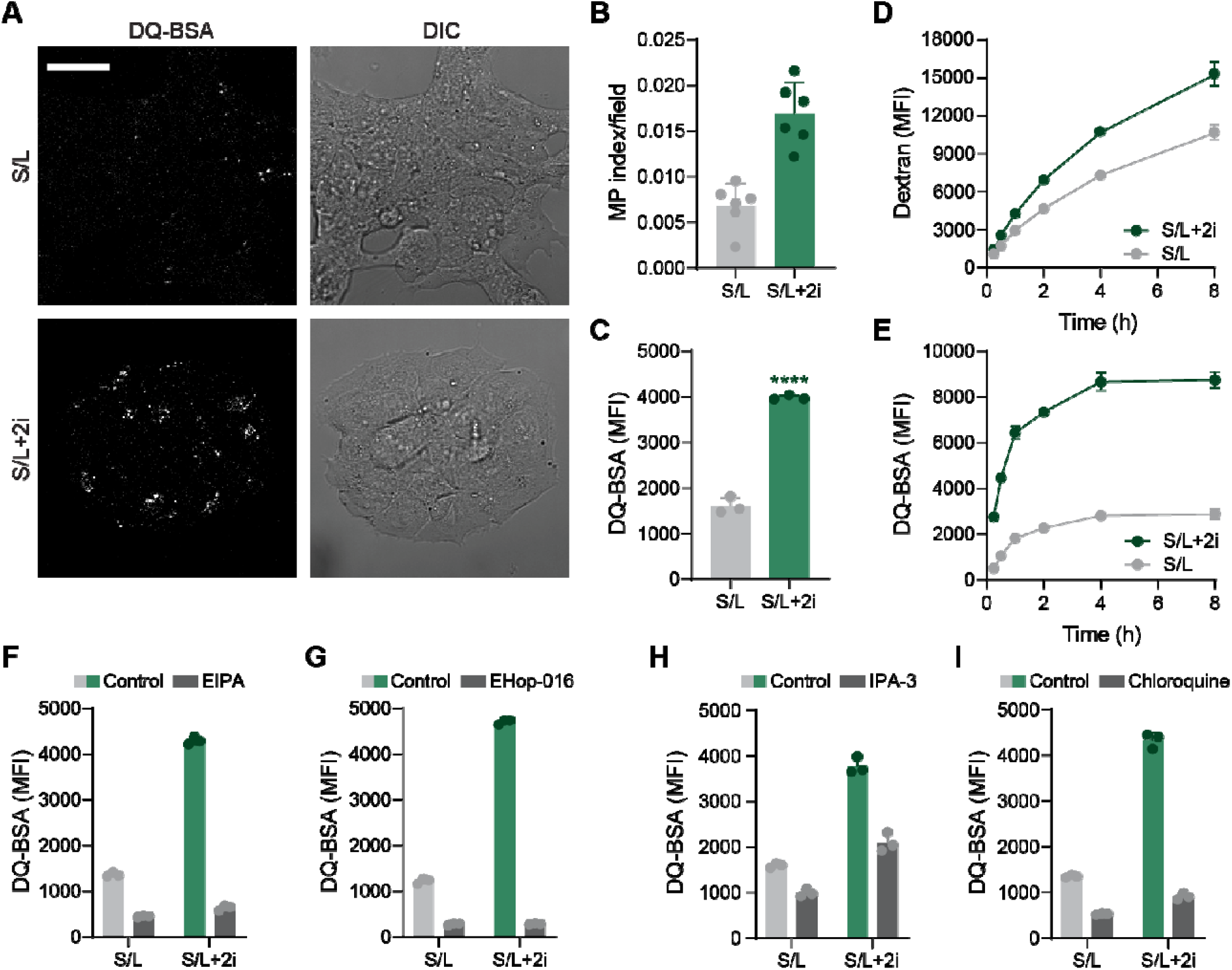
ESCs engage in constitutive macropinocytosis. (**A**) Representative fluorescent and differential interference contrast (DIC) images of S/L and S/L+2i cultured ESCs treated with DQ-BSA for 1 h. (**B**) Quantification of DQ-BSA in S/L and S/L+2i cultured ESCs. Six fields were assessed per culture condition. (**C**) Flow cytometry quantification of DQ-BSA signal in S/L and S/L+2i cultured ESCs. (**D** and **E**) Rate of Dextran (D) and DQ-BSA (E) in S/L and S/L+2i cultured ESCs. (**F**-**I**) Flow cytometry quantification of DQ-BSA signal in S/L and S/L+2i cultured ESCs following pre-treatment with the indicated inhibitors. Data are mean ± SD, n=3 independent samples (C to I). Significance was assessed by unpaired two-tailed *t* test [(C)], (\*\*\*\**p*<0.0001). Scale bar, 25 µm.

To specifically assess protein uptake, we incubated cells with a fluorescently labeled 70-kDa Dextran and found that S/L+2i-cultured ESCs took up the labeled protein at a faster rate than cells cultured in S/L (Fig. 2D, Supplemental Fig. 2A). Enhanced protein uptake was accompanied by elevated digestion, as DQ-BSA fluorescence increased more quickly and plateaued at an overall higher steady-state when cells were maintained in the presence of 2i (Fig. 2E, Supplemental Fig. 2B). This phenotype was independent of medium formulation: ESCs cultured in a serum-free medium (2i/L, which contains BSA as a protein source) exhibited similarly elevated DQ-BSA signal as S/L+2i-cultured ESCs despite differences in base media and the presence of serum (Supplemental Fig. 2C). Together, these results indicate that naïve ESCs have a higher capacity to take up and degrade extracellular proteins than their metastable counterparts. To put these results in context, we compared DQ-BSA hydrolysis in ESCs and KRAS-mutant PANC-1 cells, which sustain proliferation when amino acids are limiting if supplemented with an extracellular protein source^31^. DQ-BSA hydrolysis was significantly higher in S/L+2i-cultured ESCs than PANC-1 cells, indicating that naïve ESCs digest extracellular protein at a rate similar to highly-macropinocytotic cancer cells and compatible with continued cell proliferation (Supplemental Fig. 2D).

To confirm that macropinocytosis enables protein uptake and digestion in ESCs, we treated cells with a panel of established macropinocytosis inhibitors. EIPA (a N^+^/H^+^ exchange inhibitor)^25,32^, eHOP-016 (a Rac1 inhibitor)^33^ and IPA-3 (a Pak1 inhibitor)^34,35^ all significantly inhibited DQ-BSA hydrolysis (Fig. 2F-H). Similarly, chloroquine, an inhibitor of lysosomal acidification, also blunted DQ-BSA signal, reflecting the requirement for lysosomal digestion to liberate DQ-BSA fluorescence (Fig. 2I).

### Leucine independence is an intrinsic feature of naïve pluripotency

To determine whether macropinocytosis is a general feature of naïve pluripotency, we exploited the phenotypic heterogeneity of ESCs cultured in S/L. Within this heterogeneous population, cells with higher expression of naïve pluripotency factors such as Rex1 and Nanog more faithfully recapitulate the pre-implantation epiblast^11^. Using ESC lines with reporters for Rex1 and Nanog expression^36,37^, we found that in each case, DQ-BSA signal was significantly increased in cells with the highest expression of naïve pluripotency factors (Fig. 3A, B). To test whether increased hydrolysis of extracellular proteins corresponds to an enhanced capacity to withstand amino acid deprivation, we withdrew leucine from the culture medium for 24 h and monitored changes in cell fate. Consistent with the differential DQ-BSA signal, Nanog-low cells were specifically eliminated by leucine deprivation (Fig. 3C). Furthermore, cells remaining after leucine withdrawal exhibited increased expression of naïve pluripotency factors (*Esrrb, Klf4, Nanog, Zfp42/Rex1*) and decreased expression of genes representative of post-implantation epiblast states (*Fgf5, Oct6, Otx2, T*) (Fig. 3D). Accordingly, ESCs demonstrated increased self-renewal potential following transient leucine withdrawal, forming over 2-fold as many undifferentiated colonies compared with their control counterparts when reseeded at clonal density in leucine-replete medium (Fig. 3E). Together, these results demonstrate that reduced reliance on soluble amino acids is a selectable hallmark of cells in the naïve pluripotent state.

**Fig. 3.**
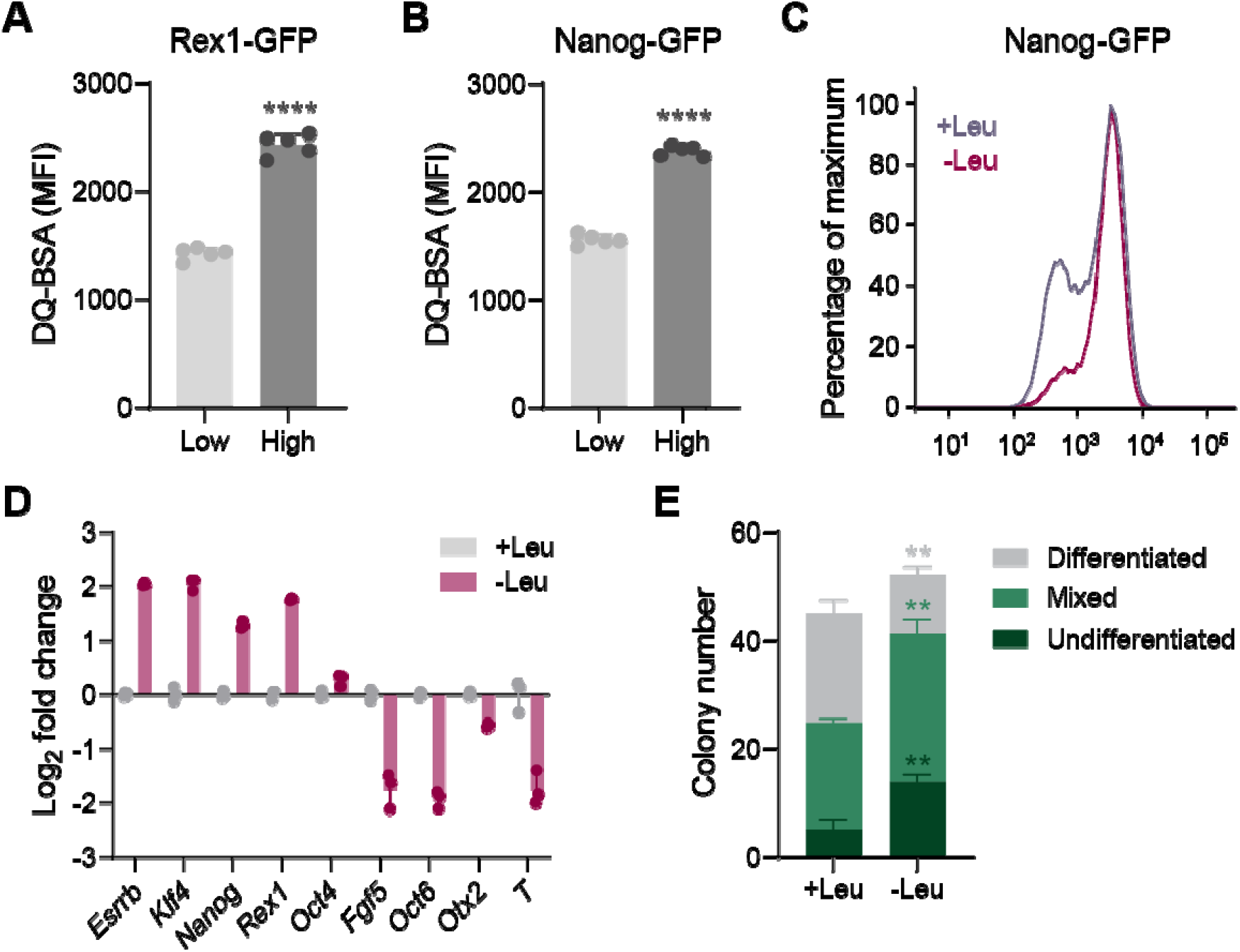
Leucine independence is a feature of naïve pluripotency. (**A** and **B**) DQ-BSA quantification in Rex1-GFP (A) and Nanog-GFP (B) S/L cultured ESCs. (**C**) Nanog-GFP expression in S/L cultured ESCs subjected to 24 h of leucine (Leu) replete or depleted conditions. (**D**) RT-qPCR of pluripotency-associated (*Esrrb, Klf4, Nanog, Zfp42/Rex1, Pou5f1/Oct4*) and early differentiation associated genes (*Fgf5, Oct6, Otx2, T*) in S/L-cultured ESCs subjected to 24 h of leucine (Leu) replete or depleted conditions. (**E**) Quantification of differentiated, mixed and undifferentiated colonies formed by ESCs seeded at clonal density in S/L medium following 24 h culture in leucine-deprived (−Leu) or control (+Leu) conditions. Data are mean ± SD, n=5 independent samples (A and B) or n=3 independent samples (D and E). Significance was assessed by unpaired two-tailed *t* test [(A), (B), and (E)] (\*\**p*<0.01, \*\*\*\**p*<0.0001).

### Lysosomal activity enables amino acid independence

We next interrogated the mechanisms enabling naïve ESCs to increase uptake and digestion of extracellular proteins. The magnitude and kinetics of DQ-BSA hydrolysis (Fig. 2E), which depends on both macromolecule uptake and digestion within the lysosome, prompted us to test whether there was an underlying difference in lysosomal activity in naïve and metastable ESCs. Staining acidic lysosomes with LysoTracker revealed that S/L+2i-cultured ESCs accumulated significantly more LysoTracker than S/L-cultured ESCs (Fig. 4A), consistent with previous reports that the KEGG lysosome gene set is the most enriched gene signature when comparing transcriptomes of 2i-cultured ESCs to S/L-cultured cells^38^. Like DQ-BSA hydrolysis, LysoTracker signal was not contingent upon 2i: S/L-cultured cells with high Nanog expression accumulated more LysoTracker than cells with low Nanog expression (Fig. 4B). These results demonstrate that lysosome accumulation and DQ-BSA hydrolysis vary in concert with cell state, rather than culture conditions.

**Fig. 4.**
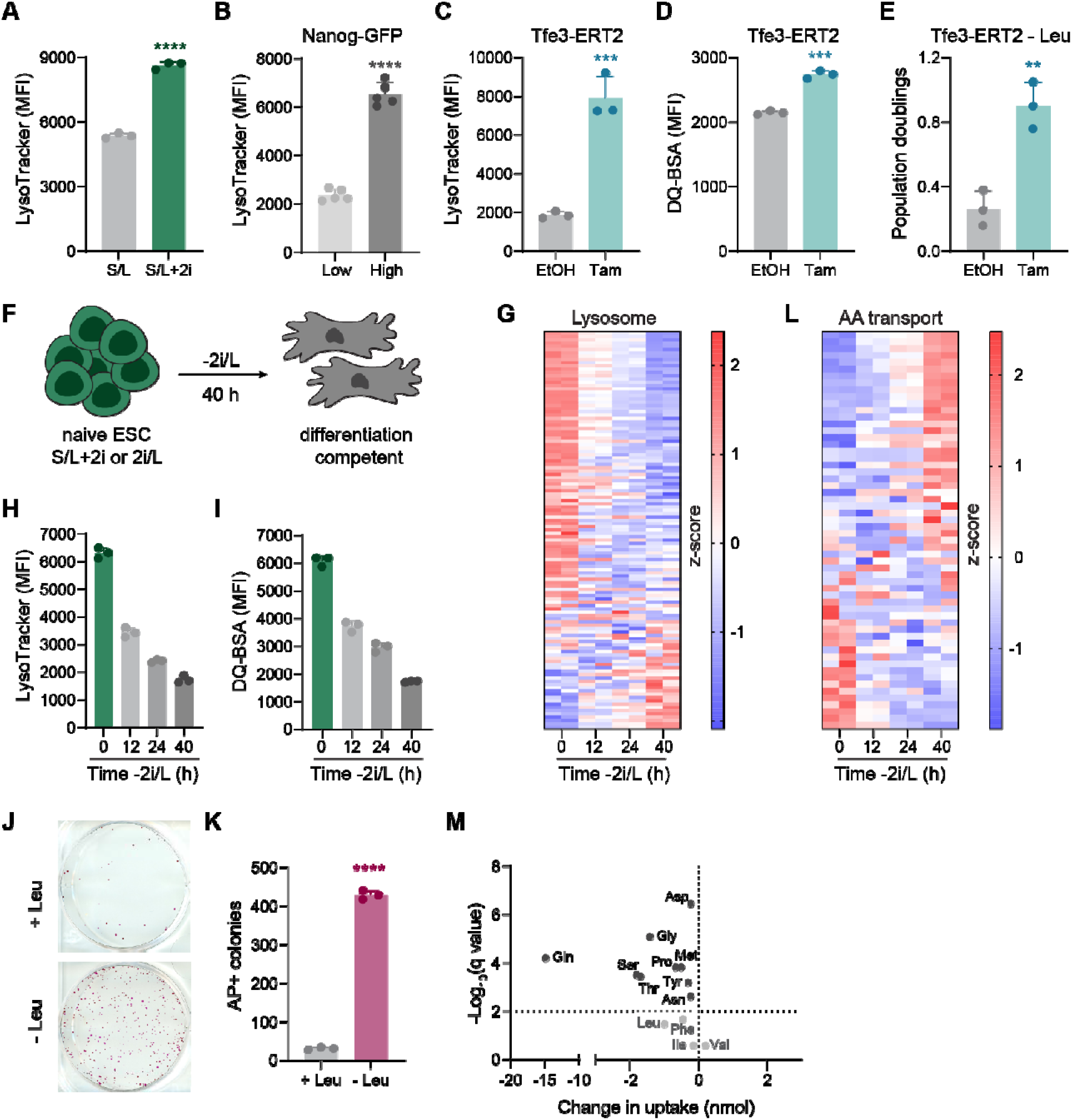
Lysosomal metabolism facilitates amino acid independence. (**A** and **B**) Quantification of LysoTracker signal in S/L and S/L+2i cultured ESCs (A) and Nanog-GFP S/L cultured ESCs (B). (**C** and **D**) LysoTracker (C) and DQ-BSA (D) signal in Tfe3-ERT2-expressing S/L cultured ESCs treated with vehicle (ethanol, EtOH) or tamoxifen (Tam). (**E**) Population doublings of Tfe3-ERT2-expressing S/L cultured ESCs treated with EtOH or Tam and subjected to leucine (Leu) withdrawal for 48 h. (**F**) Schematic of experimental setup to induce exit from naïve pluripotency. (**G**) Expression of KEGG lysosome genes in ESCs following 2i/L withdrawal for the indicated times. (**H** and **I**) Quantification of LysoTracker (H) and DQ-BSA (I) signal in 2i/L cultured ESCs subjected to 2i/L withdrawal for 12, 24 and 40 h. (**J** and **K**) Alkaline phosphatase (AP) staining of colony formation assays of S/L+2i cultured ESCs subjected to 2i/L withdrawal for 40 h in the presence or absence of leucine prior to reseeding in leucine-replete 2i/L medium to assess exit from naïve pluripotency. Representative wells are shown in J, quantification of biological replicates is shown in K. (**L**) Expression of genes encoding plasma membrane amino acid transporters in ESCs following 2i/L withdrawal for the indicated times. (**M**) Volcano plot of amino acid uptake from culture media by 2i/L cultured ESCs subjected to control or -2i/L conditions for 40 h. Media incubated with cells for the last 24 h of the experiment was assayed. Data are mean ± SD, n=3 independent samples (A, C-E, H, I, K), n=5 independent samples (B), n=2 independent samples (G and L), n=6 independent samples (M). Significance was assessed by unpaired two-tailed *t* test [(A) to (E), (K)] (\*\**p*<0.01, \*\*\**p*<0.001, \*\*\*\**p*<0.0001). Significance was assessed by multiple unpaired *t* tests and FDR was calculated using the two-stage step-up of Benjamini, Krieger, and Yekutieli [(M)].

To determine whether elevated lysosomal activity is sufficient to enable increased catabolism of exogenous protein, we introduced into S/L-cultured ESCs a version of the lysosomal biogenesis transcription factor Tfe3 fused to ERT2 (Tfe3-ERT2) to enable tamoxifen-inducible nuclear translocation and activation^39^. As expected, treating Tfe3-ERT2-expressing cells with tamoxifen for 24 h induced expression of Tfe3 target genes^39^ (Supplemental Fig. 3A) and triggered a four-fold increase in LysoTracker signal (Fig. 4C). Tamoxifen addition also significantly increased DQ-BSA hydrolysis (Fig. 4D). To test whether these changes conferred a proliferative advantage under leucine deficient conditions, we monitored changes in cell number in the presence and absence of exogenous leucine. While tamoxifen had no effect on cell proliferation in control conditions, tamoxifen treatment and Tfe3 activation was sufficient to enable cells to proliferate in leucine deficient S/L medium (Fig. 4E, Supplemental Fig. 3B). Together, these results demonstrate that heightened lysosomal macromolecular digestion provides an alternative nutrient source to sustain ESC proliferation when soluble amino acids are limiting.

### Amino acid acquisition strategies in pre- and post-implantation epiblast

In addition to its role as a transcriptional regulator of lysosome biogenesis, Tfe3 is also implicated in the maintenance of naïve pluripotency by directly binding a subset of the pluripotency transcriptional network^39^. We therefore asked whether exit from naïve pluripotency is accompanied by changes in nutrient acquisition strategies. To model exit from naïve pluripotency, we cultured ESCs in S/L+2i or 2i/L and then transitioned them to a serum-free formulation devoid of 2i and LIF (−2i/L)^16,39^ (Fig. 4F). Transcriptional profiling confirmed that in both conditions, 2i/L withdrawal induced stepwise downregulation of pluripotency genes and induction of genes associated with the early post-implantation epiblast^16^ (Supplemental Fig. 3C). Consistent with high lysosome activity defining naïve pluripotency, lysosomal genes were highly expressed in the naïve state and downregulated following 2i/L withdrawal (Fig. 4G and Supplemental Fig. 3D). Accordingly, LysoTracker signal also declined in a time-dependent fashion alongside exit from the naïve pluripotent state (Fig. 4H). These changes were accompanied by a decrease in DQ-BSA hydrolysis, indicating that the capacity to degrade extracellular protein is also reduced as cells exit the naïve pluripotent state (Fig. 4I). We therefore reasoned that withdrawing leucine alongside 2i/L would selectively eliminate cells that had exited naïve pluripotency, while selecting for cells that remained in the naïve pluripotent state, which can be assessed by returning cells to S/L+2i and quantifying their ability to form undifferentiated colonies^39^. Indeed, leucine deprivation blocked exit from naïve pluripotency, as cells subjected to leucine withdrawal retained a greater than 14-fold potential to form undifferentiated colonies when reseeded in S/L+2i (Fig. 4J).

The above results suggest that protein degradation represents a viable nutrient acquisition strategy in naïve ESCs and that cells become reliant on exogenous supplies of soluble amino acids as they dismantle naïve pluripotency. Indeed, the downregulation of lysosomal genes during exit from naïve pluripotency was accompanied by a reciprocal increase in genes encoding plasma membrane amino acid transporters (Fig. 4G, L and Supplemental Fig. 3E-G). The transcriptional upregulation of amino acid transporters was accompanied by an overall increased uptake of soluble amino acids in cells undergoing exit from naïve pluripotency compared to their naïve counterparts (Fig. 4M). Together, these data support a model in which exit from naïve pluripotency induces a metabolic switch away from digestive metabolism and towards uptake and reliance on soluble amino acids.

To put these results in the context of pre- and post-implantation embryo development, we analyzed a dataset tracking mouse development from the preimplantation blastocyst through gastrulation at single cell resolution^40^. The inner cell mass (ICM) of the embryonic day (E)3.5 pre-implantation blastocyst exhibited high lysosomal gene signatures that declined as epiblast development proceeded through pre-gastrulation at E5.5 (Fig. 5A). Reciprocally, genes encoding plasma membrane amino acid transporters were poorly expressed in the ICM and increased as the embryo progressed through implantation (Fig. 5B). Thus, early mouse development is accompanied by dynamic switches in genes related to amino acid acquisition.

**Fig. 5.**
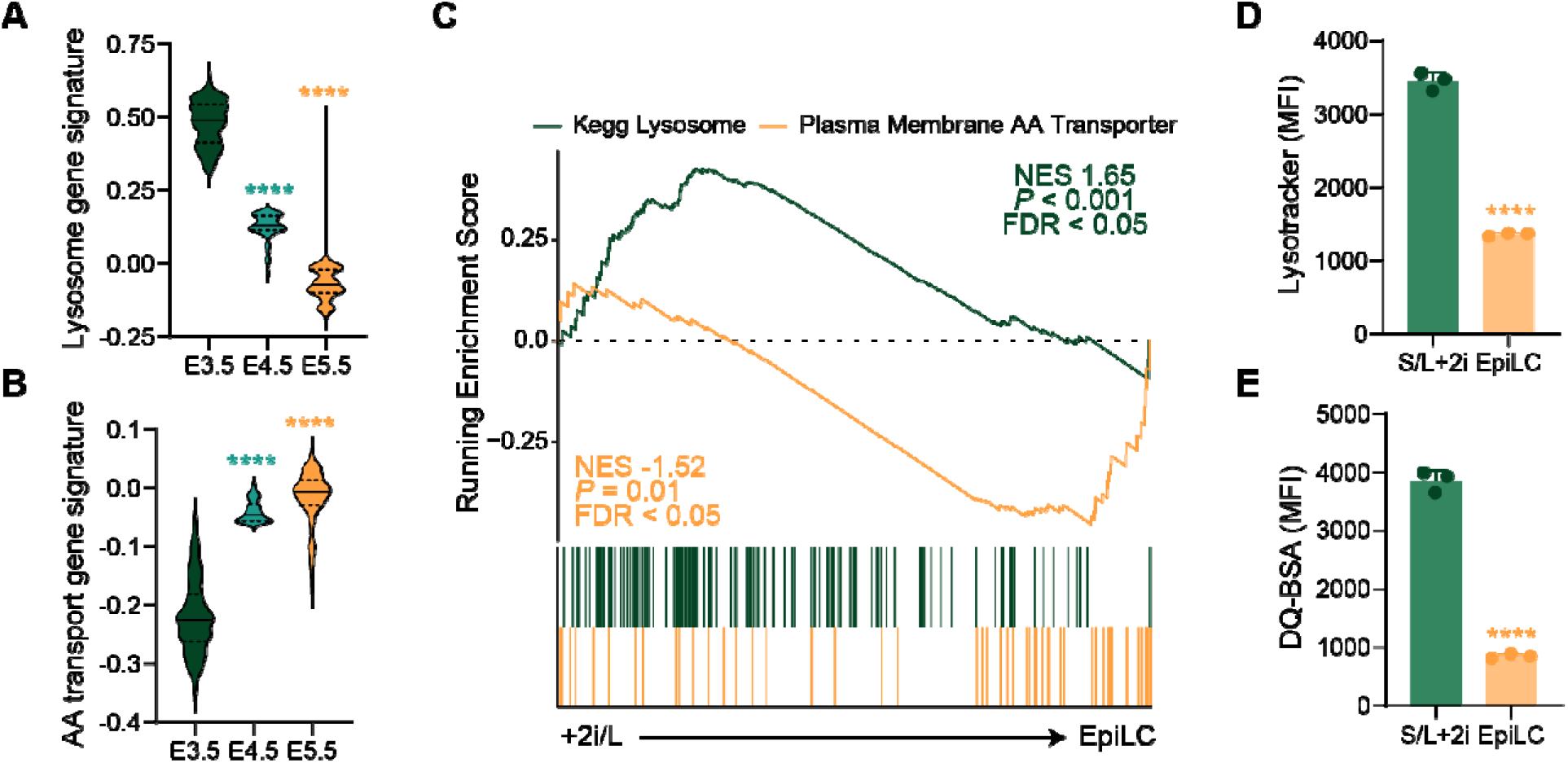
Digestive amino acid metabolism is attenuated with developmental progression. (**A** and **B**) Violin plots of scRNA-seq data ^40^ showing the expression distribution of KEGG lysosome (A) and plasma amino acid transporter (B) genes in the E3.5 ICM and E4.5 and E5.5 epiblast. (**C**) Gene set enrichment analysis of 2i/L cultured ESCs and EpiLCs ^43^ showing enrichment of KEGG Lysosome genes among genes upregulated in 2i/L and plasma membrane amino acid transporter genes enriched in the EpiLC state(**C**) Gene set enrichment analysis of 2i/L cultured ESCs and EpiLCs ^43^ showing enrichment of KEGG Lysosome genes among genes upregulated in 2i/L and plasma membrane amino acid transporter genes enriched in genes upregulated in the EpiLC state. (**D** and **E**) Quantification of LysoTracker (D) and DQ-BSA (E) in S/L+2i cultured ESCs and EpiLCs. Data are mean ± SD, n=3 independent samples (D and E). Significance was assessed in comparison to E3.5 cells by one-way ANOVA with Sidak’s multiple comparisons post-test (\*\*\*\**p*<0.0001) [(A) and (B)]. Significance was assessed by unpaired two-tailed *t* test [(D) and (E)], (\*\*\**p*<0.001, \*\*\*\**p*<0.0001).

The reciprocal regulation of genes encoding lysosomal and plasma membrane amino acid transporters was not unique to developing embryos but was also recapitulated in cell culture models of pre- and post-implantation epiblast. Naïve ESCs can be induced to adopt an epiblast-like (EpiLC) cell identity corresponding to the pre-gastrulation, post-implantation epiblast by culture with FGF2 and Activin A^14,41,42^ (Supplemental Fig. 5A, B). Comparing transcriptional profiles of ESCs and EpiLCs^43^ revealed that lysosomal genes are highly enriched among genes upregulated in naïve ESCs, whereas plasma membrane amino acid transporters are overrepresented in genes upregulated in EpiLCs (Fig. 5C). Consistent with this transcriptional switch, we found that EpiLCs exhibit sharply reduced LysoTracker and DQ-BSA signal compared to naïve ESCs (Fig. 5 D, E). Together, these results suggest that epiblast development is associated with metabolic reprogramming from digestive to absorptive nutrient acquisition strategies.

#### Mouse preimplantation blastocysts engage in constitutive macropinocytosis

The high lysosomal gene signatures in the pre-implantation epiblast (Fig. 5A) led us to ask whether the capacity to degrade extracellular protein is a feature of pre-implantation development. To address this question, we incubated E3.75 blastocyst stage mouse embryos with DQ-BSA (Fig. 6A). Within 30 min of treatment, intact blastocysts showed robust DQ-BSA signal localized to the outermost trophectoderm layer (Fig. 6B). This signal was eliminated by EIPA, indicating that trophoblast cells are capable of acquiring macromolecules via macropinocytosis and delivering them to the lysosome for digestion.

**Fig. 6.**
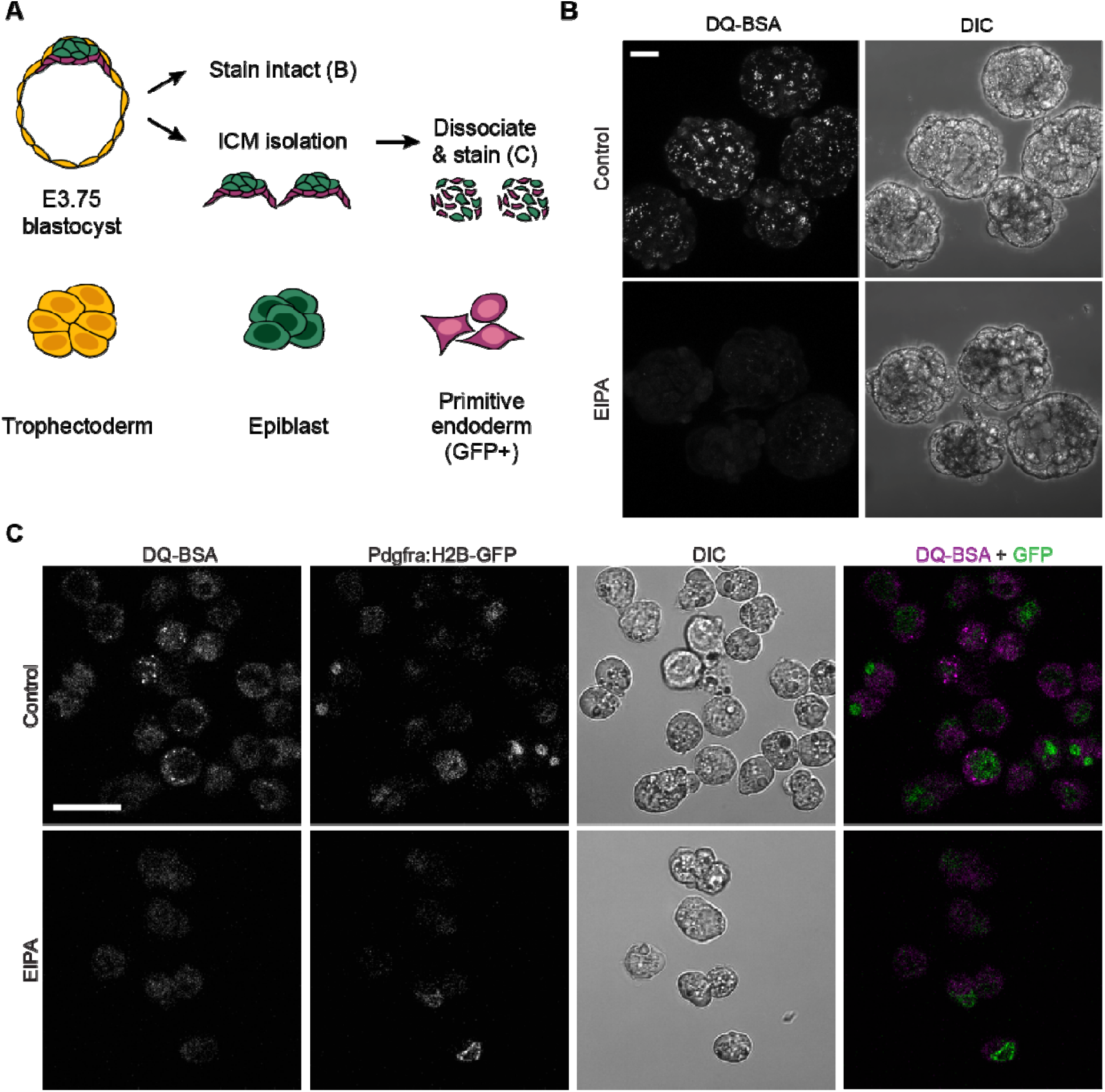
Cells in pre-implantation blastocyst engage in macropinocytosis. (**A**) Schematic of experimental workflow in E3.75 mouse blastocysts. (**B**) Representative fluorescent and differential interference contrast (DIC) images of control and EIPA pre-treated E3.75 mouse blastocysts incubated with DQ-BSA. (**C**) Representative fluorescent and DIC images of control and EIPA pre-treated ICM dissociated from E3.75 *Pdgfra*^*H2B-GFP/+*^ blastocysts and incubated with DQ-BSA. GFP demarcates primitive endoderm. Scale bars, 25 µm.

The trophectoderm layer encapsulates the ICM, which is comprised of pluripotent epiblast (EPI) and extra-embryonic primitive endoderm (PrE) lineages. To determine whether these lineages engage in macropinocytosis, we immunosurgically isolated and dissociated the ICMs from E3.75 *Pdgfra*^*H2B-GFP/+*^ blastocysts^44,45^. Here, Pdgfra-driven GFP expression marks the PrE, enabling us to identify PrE and/or EPI cells and determine whether they exhibit constitutive protein uptake and degradation. Indeed, both GFP-positive (PrE) and GFP-negative (EPI) cells internalized and hydrolyzed DQ-BSA in an EIPA-dependent manner (Fig. 6C). Together, these results reveal that all three lineages of the pre-implantation embryo engage in constitutive macropinocytosis.

## Discussion

Here we show that the pre- and post-implantation EPI states *in vitro* and *in vivo* are marked by a fundamental switch in nutrient acquisition strategies. We find that pluripotent stem cells representing the pre-implantation epiblast effectively degrade protein to sustain viability without uptake of soluble amino acids while post-implantation epiblast-like cells enhance uptake of soluble amino acids and are thus dependent upon exogenous amino acids for viability and proliferation. These results provide an explanation for the long-standing observation that mammalian embryos can develop in the absence of soluble amino acids^3-5^. Furthermore, our results are consistent with an array of studies using human, mouse and cow embryos which demonstrated that amino acid uptake and release is inversely correlated with blastocyst development: in other words, the healthiest embryos exhibited the lowest turnover of exogenous soluble amino acids^46-48^. Taken together, these results support a model in which pre-implantation development is characterized by digestive amino acid acquisition strategies that are traded upon implantation for more conventional, absorptive amino acid acquisition strategies.

Our finding that macropinocytosis is a constitutive feature of pre-implantation embryo development raises the possibility that digestion of both external and locally produced protein (for example, from proteins deposited in the oocyte or released from dying cells within the blastocyst) provides a meaningful source of amino acids for developing embryos. Suggestively, epithelial phagocytosis of apoptotic corpses has recently been implicated in error correction in the early embryo^49^, and this process would conceivably provide a local source of exogenous protein for embryo catabolism. Indeed, mouse embryos can develop in the complete absence of any fixed nitrogen, although development proceeds more readily in the presence of a protein source such as BSA^5^. The embryo may derive several advantages from using macromolecules as nutrient sources: low expression of plasma membrane amino acid transporters may safeguard against excessive uptake of nutrients that could fuel oxidative metabolic pathways and concomitant production of damaging reactive oxygen species, and reduced transporter expression insulates embryos from environmental nutrient fluctuations prior to implantation.

Consistently, mouse embryos only need 4 cations (Na^+^, K^+^ Ca^2+^, Mg^2+^), 3 anions (Cl^-^, PO_4_^3-^, HCO_3_^-^), pyruvate and water to develop to the blastocyst stage^50^. In this manner, mammalian development bears some conceptual similarity to oviparous development in which many of the nutritional needs of early development can be met autonomously.

Macropinocytosis may provide additional benefits beyond nutrient acquisition to developing embryos. For example, macropinocytosis could facilitate recycling nutrients from debris of dying cells that are a feature of normal development. Macropinocytosis also involves significant membrane remodeling which can enable plasma membrane repair^51^. Plasma membrane turnover can also have significant consequences for cellular signaling cascades and may provide a mechanism for pre-implantation blastocysts to shield themselves from inductive cues prior to implantation^52,53^. While the precise signals linking cell states to macropinocytosis induction remain to be elucidated, our data indicate that enhanced lysosomal capacity facilitates nutrient acquisition from exogenous macromolecules (Fig. 4). Given that the transcription factors known to induce lysosomal gene expression can also directly bind *cis*-regulatory elements of genes controlling stem cell identity^39,54^, our results raise the possibility that adult stem cell compartments may likewise harbor capacity to digest exogenous macromolecules as fuel sources. In this scenario, efforts to target or exploit macropinocytosis as anti-cancer therapy^55^ may need to consider potential effects on stem cell homeostasis.

More broadly, our results underscore the diversity of metabolic strategies that can support rapid cell division. To date, most studies on metabolism in proliferating cells have used cancer cells or immune cells as model systems. A common theme in these models is the requirement for elevated uptake of a variety of nutrients to support cell growth^7,56^. Here, our findings that ESCs sustain rapid proliferation even without soluble essential amino acids supports the notion that metabolic strategies that enable cell division can vary widely between cell types and that continued study of proliferative metabolism in distinct conditions and cell states is likely to uncover unappreciated diversity in the metabolic networks that support cell growth. Indeed, emerging evidence indicates that the specification and differentiation of early embryonic lineages is likely accompanied by significant metabolic rearrangements that induce lineage-or state-specific dependencies. For example, glutamine is dispensable for the developing ectoderm but required for mesoderm and endoderm specification; similarly, lysosomal transcriptional networks are specifically required for endoderm specification^57,58^. Further understanding of metabolic requirements that characterize developmental stages in different cell lineages at distinct developmental stages will likely yield critical insight into the etiology of developmental disorders such as inborn errors of metabolism as well as provide guidance for interventions to safeguard normal development.

## Acknowledgements

We thank members of the Finley lab for helpful discussion, Bryan King for technical guidance, and the Media Preparation Facility at MSKCC for specialized media production. B.T.J. is an NICHD Ruth L. Kirschstein Predoctoral fellow (1F30HD107943-01) and is supported by a Medical Scientist Training Program grant from the NIGMS of the National Institutes of Health under award number T32GM007739 to the Weill Cornell/Rockefeller/Sloan Kettering Tri-Institutional MD-PhD Program. K.I.P. was supported by a Bruce Charles Forbes Pre-Doctoral Fellowship (MSKCC). L.W.S.F. was a Searle Scholar. This research was additionally supported by the Memorial Sloan Kettering Cancer Center Support Grant P30CA008748, grants to A.K.H. from the NIH (R01DK127821 and R01HD094868), and grants to L.W.S.F. from the Starr Foundation (I12-0051), the Tri-Institutional Stem Cell Initiative (2019-007), and the Basic Research Innovation Award from the Sloan Kettering Institute.

## Competing interests

The authors declare no conflicts of interest.

## Methods

### Cell culture

Mouse ESCs (ESC-1, ESC-2) were previously generated from C57BL/6□×□129S4/SvJae F_1_ male embryos^18^ ESC-1 cells were used for all experiments unless otherwise stated. Nanog-GFP reporter ES cells were a gift from R. Jaenisch^36^. *Rex1::GFPd2* cells were a gift from A. Smith^37^. ESCs were maintained on gelatin-coated plates in the following maintenance media: serum/LIF (S/L), serum/LIF+2i (S/L+2i) or 2i/LIF (2i/L). S/L medium contained Knockout DMEM (10829018; Thermo Fisher Scientific) supplemented with 10%□fetal bovine serum (FBS; Gemini), 0.1□mM 2-mercaptoethanol, 2□mM L-glutamine and 1,000□U/mL LIF (Gemini). S/L+2i contained S/L medium supplemented with an additional 3□µM CHIR99021 (Stemgent) and 1□µM PD0325901 (Stemgent) (2i). 2i/L medium consisted of a 1:1 mix of DMEM/F-12 (11320033; Gibco) or DMEM (11960069; Gibco) and Neurobasal medium (21103049; Gibco) with N-2 supplement (1:100 17502048; Gibco), B-27 supplement (1:100 17504044; Gibco), 0.1 mM 2-mercaptoethanol, 2□mM L-glutamine, LIF and 2i. S/L cells were converted to S/L+2i or 2i/L over three passages and adapted cells were maintained for a maximum of nine passages.

To induce exit from naïve pluripotency, ESCs cultured in either S/L+2i or 2i/L were seeded at least 12□h before washing with PBS and changing into 1:1 mix of glutamine-free DMEM/F12 or glutamine-free DMEM and Neurobasal medium, 1:100 N-2 supplement, 1:100 B-27 supplement, 2 mM L-glutamine, and 0.1 mM 2-mercaptoethanol at the indicated time before collection (12, 24 or 40□h without LIF and 2i). When leucine withdrawal was performed while cells were undergoing exit, in-house produced leucine-free DMEM and Neurobasal media were used.

EpiLC cells were derived from ESCs using a previously described protocol^59^ with slight modifications. ESCs cultured in S/L+2i were seeded onto Fibronectin-coated plates and cultured in medium containing a 1:1 mix of DMEM-F12 and Neurobasal medium supplemented with N2 (1:200), B-27 (1:100), 2 mM L-glutamine, 0.1 mM 2-mercaptoethanol, 12 ng/mL FGF2, 20 ng/mL Activin A and 1% Knockout serum replacement. All EpiLC cell experiments were performed 48 h after switching to differentiation medium. PANC-1 cells were obtained from ATCC and maintained in DMEM (produced in house) supplemented with 10% FBS (Gemini).

### Growth curves and biomass accumulation

ESCs maintained in S/L or adapted to S/L+2i were seeded at a density of 40,000 cells/well in 12-well plates. After 24 h, three wells of each cell line were counted to determine starting cell number and biomass. The remaining wells were washed with PBS and changed to medium containing 1:1 mix of glutamine-, leucine-, methionine-, lysine-or threonine-free DMEM and Neurobasal media (produced in house), 10% dialyzed FBS (Gemini), 0.1 mM 2-mercaptoethanol, 2 mM L-glutamine (except for the glutamine withdrawal condition) and 1,000 U/mL LIF (Gemini). This medium was further supplemented with 3µM CHIR (Stemgent) and 1µM PD0325901 (Stemgent) for the S/L+2i conditions. For the amino acid replete arm of each growth curve, the corresponding amino acid was added back to deficient media. Cells were harvested and counted 48 h later using a Beckman Coulter Multisizer 4e gated at 400-10,000 fL.

Final cell counts were normalized to starting cell number. Biomass volume was calculated by multiplying cell numbers by the median cell volume obtained from the Beckman Coulter Multisizer 4e measurement. Final biomass was normalized to biomass at the onset of amino acid withdrawal.

### Viability assay

S/L- and S/L+2i-adapted ESCs were seeded at a density of 24,000 cells per well of a 24-well plate in triplicate. At the indicated timepoints, the cells were washed with PBS and changed to medium containing a 1:1 mix of glutamine- and leucine-free DMEM and Neurobasal media, 10% dialyzed FBS, 2 mM L-glutamine, 0.1 mM 2-mercaptoethanol and 1,000 U/mL LIF with or without leucine and 2i. 48 h after seeding, all cells were collected with trypsin, pelleted, and resuspended in flow buffer (PBS with 2% FBS, 0.5 mM EDTA and 0.05% sodium azide) containing Propidium iodide (PI, 2.5 µg/mL; Biotium). PI incorporation was measured on a LSRFortessa flow cytometer using FACSDiva software v.8.0 (BD Biosciences). Analysis of propidium iodide exclusion was performed using FCS Express v.7.12.0007.

### Extracellular protein uptake and hydrolysis

Cells were seeded at a density of 24,000 cells per well of a 24-well plate in experiment-specific medium conditions and changed into fresh media the next day. After 24 h, cells were supplemented with intracellular dyes (20 µg/mL DQ Green or DQ Red BSA (Thermo Fisher Scientific), 250 µg/mL Dextran Texas Red 70,000 MW Neutral (Thermo Fisher Scientific), 100 nM LysoTracker Red DND-99 (Thermo Fisher Scientific)); and inhibitors (50 µM 5-(N-ethyl-N-isopropyl)-Amiloride (EIPA; Fisher Scientific), 8 µM EHop-016 (Selleck Chemicals), 10 µM IPA-3 (Fisher Scientific) or 50 µM chloroquine (Sigma Aldrich)). Cells were incubated with dyes for 1 h with or without 1 h pre-treatment with indicated inhibitors. For the DQ-BSA and Dextran time-course experiments, dyes were added at the indicated time points prior to harvest. Cells were washed with PBS twice and resuspended in 4’,6 Diamidino 2 Phenylindole, Dihydrochloride (DAPI; Thermo Fisher Scientific) containing flow cytometry buffer (PBS with 2% FBS, 0.5 mM EDTA and 0.05% sodium azide). DQ-BSA, Dextran and LysoTracker incorporation was measured on the LSRFortessa flow cytometer using FACSDiva software v.8.0 (BD Biosciences) and analyzed using FCS Express v.7.12.0007.

Nanog-GFP ESCs cultured in S/L were treated with LysoTracker or DQ-BSA (100 nM and 20 µg/mL respectively) for 1 h and washed with PBS twice before analysis. LysoTracker and DQ-BSA signal in GFP-high (top 5%) and GFP-low (bottom 5%) populations were compared. For leucine-deprivation experiments, cells were seeded in S/L medium and 24 h later were washed with PBS and changed to leucine-free DMEM/Neurobasal S/L medium with or without leucine. After 24 h, cells were harvested and GFP signal was measured on the LSRFortessa flow cytometer using FACSDiva software v.8.0 (BD Biosciences) and analyzed using FlowJo v.10.8.1.

### Colony formation assay

ESCs maintained in S/L conditions were seeded at a density of 150,000 cells per well of a 6-well plate (3 wells per experimental group). The following day cells were washed with PBS and changed to leucine-free medium containing 1:1 mix of glutamine- and leucine-free DMEM and Neurobasal media, 10% dialyzed FBS, 2 mM L-glutamine, 0.1 mM 2-mercaptoethanol and 1,000 U/mL LIF with or without leucine. After 24 h cells were counted and reseeded in technical triplicate at a density of 500 cells per well in leucine-replete S/L medium. Medium was refreshed 3 days after seeding and 3 days later cells were fixed with citrate/acetone/3% formaldehyde and stained with Leukocyte Alkaline Phosphatase Kit (Sigma-Aldrich) according to the manufacturer’s instructions. Colonies were classified as differentiated, mixed, and undifferentiated by a blinded investigator. The mean of three technical triplicates is reported as a single datapoint.

To assess exit from naïve pluripotency, ESCs maintained in S/L+2i were subjected to 40 h of 2i and LIF withdrawal in serum-free medium (1:1 mix of glutamine- and leucine-free DMEM and Neurobasal media, 1:100 N-2 supplement, 1:100 B-27 supplement, 2 mM L-glutamine, 0.1 mM 2-mercaptoethanol) with or without leucine in biological triplicates. After 40 h cells were counted and reseeded at a density of 2,000 cells per well in maintenance S/L+2i medium. Each biological replicate was seeded in technical triplicates. Colonies were stained with alkaline phosphatase as described above and quantified using ImageJ’s particle analysis tool by a non-blinded investigator. The mean of three technical triplicates is reported as a single datapoint.

### Generation of a Tfe3-ERT2 ESC line

ESC-1 cells were transfected with a tamoxifen-responsive version of murine Tfe3 using the Piggybac system. pPB-CAG-Tfe3::ERT2-pgk-hph was a gift from Austin Smith (Addgene plasmid # 48756; http://n2t.net/addgene:48756). Cells were transfected with the Piggybac plasmid plus transposase at a 3:1 ratio using Fugene HD (Promega) and selected with G418 (300 µg/mL, Gibco). Tfe3 nuclear translocation was induced by treating transfected cells with 4-hydroxytamoxifen (1µg/mL, Sigma Aldrich).

### Quantification of gene expression

RNA was isolated using TRIzol (Invitrogen) according to manufacturer’s instructions and 200 ng RNA was used for cDNA synthesis using the iScript cDNA Synthesis kit (Bio-Rad). Quantitative real-time PCR was performed in technical triplicates using Power SYBR Green Master Mix (Thermo Fisher Scientific) and analyzed on QuantStudio 5 (Applied Biosystems). Data were generated using RNA from 3 independent wells for each condition. *Actin* was used as endogenous control. A list of all primer sequences is provided in Supplementary Table 1.

### RNA-seq analysis

RNA was isolated with TRIzol and quantified with a Qubit 3.0 fluorometer (Thermo Fisher Scientific). RNA-seq libraries were generated using the TruSeq Stranded mRNA Library Prep Kit (20020594, Illumina), pooled and sequenced at the Memorial Sloan Kettering Cancer Center Integrated Genomics Operation. fastp^60^ was used to filter and trim the libraries; mapping was performed using STAR aligner against the mm10 mouse genome assembly using default parameters. Gene counts were calculated using featureCounts^61^ and inputed into DESeq2^62^ for quality control analysis, size normalization and variance dispersion corrections. For heatmaps, log_2_ normalized counts were generated by DESeq2 for z-score calculation. Z-scores were calculated as z = (x-µ)/*σ*, where µ and *σ* are the mean expression and standard deviation across all samples for a given gene. For regression analysis, normalized gene scores for each sample were calculated by subtracting the average log_2_ normalized counts of a given gene across all samples from the log_2_ normalized count of each gene. All genes from all samples were plotted using ggplot2^63^ and the regression line was calculated using the geom_smooth function with method=‘lm’.

### Nutrient consumption

ESC-1 cells cultured in 2i/L medium were seeded at a density of 90,000 cells/well in 6-well plates in 2i/L medium or induced to exit pluripotency (−2i/L) for 40 h as described above. 24 h before harvest, medium was aspirated and cells were given 1 mL of fresh medium. A 6-well plate with no cells was incubated in parallel with 1 mL medium as a media-only control for normalization. 24 h later, media samples were collected and spun down at 1,700 x *g* for 5 min in a pre-chilled centrifuge. 50 µL of supernatant per sample was mixed with 1□mL of ice-cold 80% methanol supplemented with 2□μM deuterated 2-hydroxyglutarate (*D*-2-hydroxyglutaric-2,3,3,4,4-d_5_ acid (d5-2HG)) as an internal standard. Samples were incubated overnight at -20°C. Samples were spun down at 20,000 x *g* for 20 min at 4°C and supernatants were dried in an evaporator (Genevac EZ-2 Elite) for further derivatization for GC-MS. First, 50 µL of 40□mg□mL^−1^ methoxyamine hydrochloride in pyridine was added and the samples were incubated at 30°C for 2 h with shaking. Next, 80□μL of *N*-methyl-*N*-(trimethylsilyl) trifluoroacetamide□(Thermo Fisher Scientific) and 70□μL ethyl acetate (Sigma-Aldrich) were added, followed by a 30 min incubation at 37□°C. Metabolites were analyzed on an Agilent 7890A gas chromatograph coupled to an Agilent 5977C mass selective detector. The gas chromatograph was obtained using splitless injection mode with constant helium gas flow at 1□mL□min^−1^; 1□μL of derivatized metabolites was injected onto an HP-5ms column and the gas chromatograph oven temperature ramped up from 60□°C to 290°C over 25□min. Peaks representing compounds of interest were analyzed using MassHunter v.B.08 (Agilent Technologies) and then normalized to the internal standard (d5-2HG) peak area by quantifying the following ions: d5-2HG, 354□*m*/*z*; asparagine, 231 *m*/*z*; aspartate, 334 *m*/*z*; glutamine, 362 *m*/*z*; glycine, 276 *m*/*z*; isoleucine, 260□*m*/*z*; leucine, 260□*m*/*z*; methionine, 250□*m*/*z*; phenylalanine, 294 *m*/*z*; proline, 216 *m*/*z*; serine, 306 *m*/*z*; threonine, 320□*m*/*z*; tyrosine, 382 *m*/*z*; valine, 246 *m*/*z*. d5-2HG-normalized peak area of metabolites from each sample was further normalized to control (media-only) normalized peak areas to determine the fraction of each amino acid that was consumed. Values were further normalized to starting amino acid abundance to determine the amount of amino acid consumed in each sample.

### Gene expression correlation

RNAseq of naïve ESCs or EpiLCs was obtained from previously published data^43^. Genes were ranked by log_2_ fold change of 2i/L relative to EpiLC, and gene set enrichment analysis^64^ of the gene set KEGG lysosome (KEGG_LYSOSOME, M11266) or a custom plasma membrane amino acid transporter gene set (Supplementary Table 2) was performed and graphed using clusterProfiler v4.0^65^.

### Single-cell RNA-seq analysis

scRNA-seq data from mouse E3.5-E5.5 embryos were obtained from endoderm-explorer.com^40^. Genes detected in fewer than 10 cells were filtered out. The count matrix was normalized for library size, multiplied by a high value (10,000) and log transformed with a pseudo-count of 0.1 using scprep (https://github.com/KrishnaswamyLab/scprep) as previously described ^40^. This count matrix was used for MAGIC gene expression imputation^66^ with default parameters. To calculate lysosomal and plasma membrane amino acid transporter gene signatures, MAGIC-imputed data was loaded into Seurat^67^ and the AddModuleScore function was performed with the gene set KEGG lysosome (KEGG_LYSOSOME, M11266) or a custom plasma membrane amino acid transporter gene set (Supplementary Table 2). For E3.5, cells annotated as ICM, EPI, or PrE were included, while for E4.5 and E5.5 only EPI-annotated cells were used for gene signature analysis.

### Immunocytochemistry

ESCs were seeded on gelatin coated plastic coverslips (Ibidi) and medium was changed 24 h later. Cells were allowed to grow for additional 24 h before treatment with 100 µg/mL DQ Green BSA (Thermo Fisher Scientific) for 1 h. Cells were then washed twice in PBS and imaged using a Zeiss Axio Vert.A1 inverted microscope with a black and white camera (Axiocam MRm). Images were analyzed using ImageJ v1.53t. Macropinocytosis index per field was calculated as previously described^68^.

### Embryo imaging

Embryonic days (E) were determined by considering noon of the day of detection of the copulation vaginal plug as E0.5. Preimplantation embryos corresponding to E3.75 were flushed from uterine horns of pregnant females and washed in Flushing and Holding Media (FHM, EMD Millipore). Zona pellucidae were removed by incubation in Acid Tyrode’s solution (EMD Millipore) and denuded blastocysts were washed in FHM prior to staining (whole embryo staining) or immunosurgery (ICM staining)^69^. To isolate individual ICM cells, blastocysts were incubated in 1:1 mix of anti-mouse lymphocyte serum (Cedarlane) and FHM and incubated at 37°C for 15 min. Blastocysts were then washed well in FHM and incubated in guinea pig complement (Cedarlane) for an additional 8-10 min at 37°C. Lysed trophectoderm cells were removed by trituration with pulled glass capillaries (Sutter Instruments). Isolated ICM were dissociated into single cells with 0.5% Trypsin-EDTA, followed by mechanical disruption with pulled capillaries of serially smaller bore sizes.

Intact blastocysts or dissociated ICM cells were treated with EIPA or vehicle and incubated for 30 min in a humidified incubator at 37°C and 5% CO_2_. Samples were then treated with DQ-BSA and returned to the incubator for an additional 30 min. DQ-BSA treated blastocysts or cells were washed through 4-5 drops of EmbryoMax KSOM Mouse Embryo Medium or PBS supplemented with 10% FBS solution, respectively. Samples were stained with DAPI and transferred to microdrops of 10% FBS solution on 35 mm glass bottom dishes (MatTek) for imaging. Images were acquired on a Zeiss LSM 880 laser-scanning confocal microscope using an EC Plan-Neofluar 40×/NA1.30 oil immersion objective at 1-μm z-intervals. Fluorescence was excited using a 405-nm diode (Hoechst 3342), 488-nm argon, and 561-nm DPSS-561-10.

### Mouse strains and husbandry

All animal work was approved by Memorial Sloan Kettering Cancer Center’s Institutional Animal Care and Use Committee (protocol 03-12-017, Hadjantonakis PI). Animals were housed in a pathogen-free facility under a 12-hour light cycle. All embryos used for this study were obtained from natural matings of virgin females of 5-10 weeks of age. Mouse strains used in this study were: *Pdgfra*^*H2B-GFP*^ (Jackson Labs, Bar Harbor, ME, USA/stock ID: 007669)^44^, and wild-type CD1 (Charles River)

### Statistics and reproducibility

Prism 9 (GraphPad) software was used for statistical analyses, except for GSEA and RNA-seq analysis which were performed in R v4.2. Further sc-RNAseq processing was performed using Python v3.8. Statistical tests and *P* values are reported in the figure legends. Experiments were performed in biological triplicate or as indicated in the figure legends and were repeated two or more times.

## Data code and availability

RNAseq data supporting the findings of this study have been deposited in the Gene Expression Omnibus under the accession code GSE216356 (reviewer token: cpupowekpjwdrkt). Alignment was performed against the mouse mm10 genome assembly. EpiLC conversion RNA-seq was obtained from previously published data^43^. E3.5-E5.5 scRNA-seq data was obtained from endoderm-explorer.com^40^.

---

**Supplemental Fig.1.**
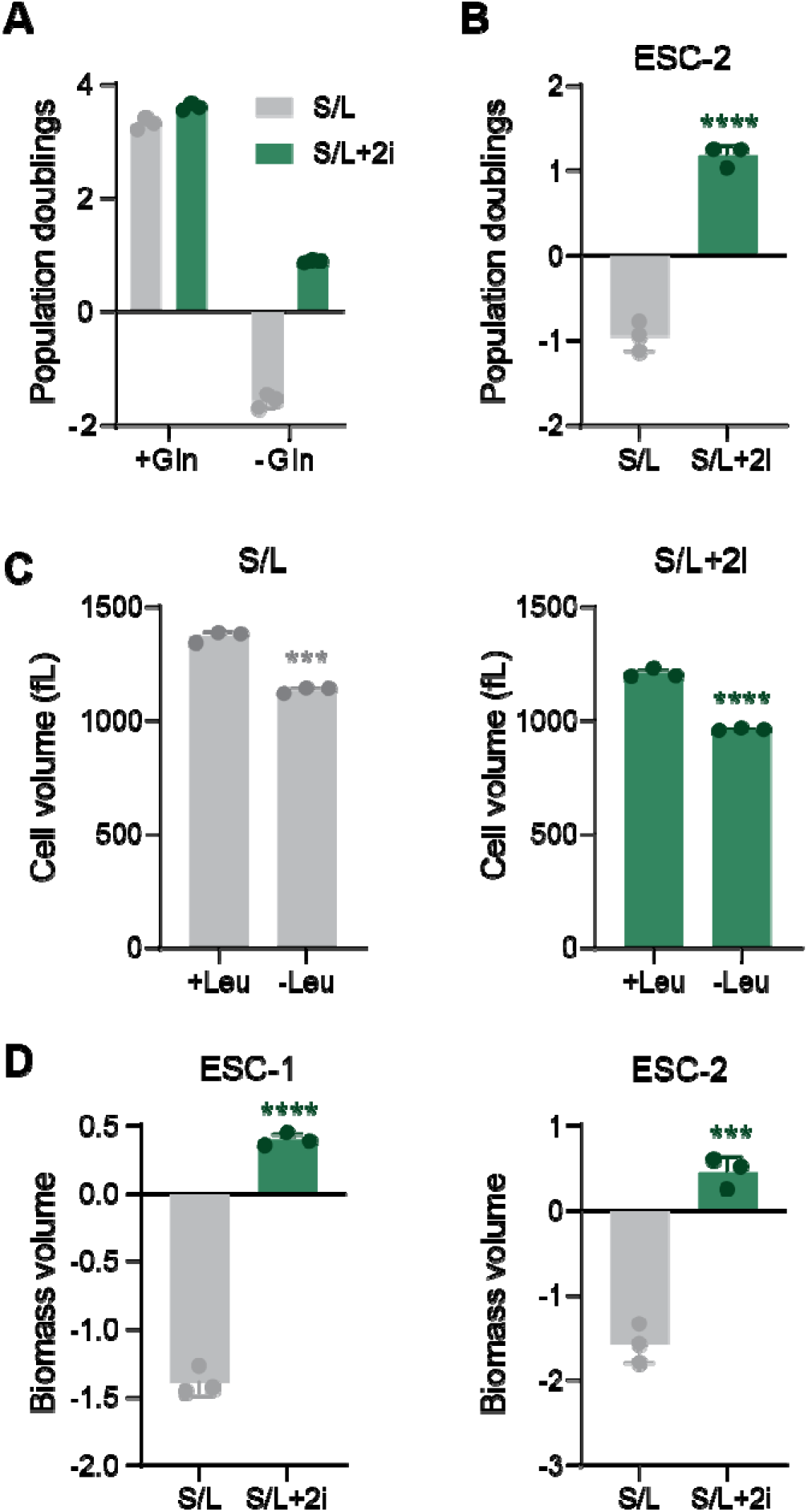
Naïve ESCs grow without exogenous leucine. (**A**) Population doublings of S/L and S/L+2i cultured ESCs subjected to glutamine (Gln) replete or depleted conditions for 48 h. (**B**) Population doublings of S/L and S/L+2i cultured ESCs (ESC-2) subjected to leucine (Leu) depleted conditions for 48 h. (**C**) Cell volume of S/L and S/L+2i cultured ESCs subjected to leucine (Leu) replete or depleted conditions for 48 h. (**D**) Change in biomass of S/L and S/L+2i cultured ESCs (ESC-1 and ESC-2) subjected to leucine (Leu) depleted conditions for 48 h. Data are mean ± SD, n=3 independent samples. Significance was assessed by unpaired two-tailed *t* test [(C), (D)] (\*\*\**p*<0.001, \*\*\*\**p*<0.0001).

**Supplemental Fig.2.**
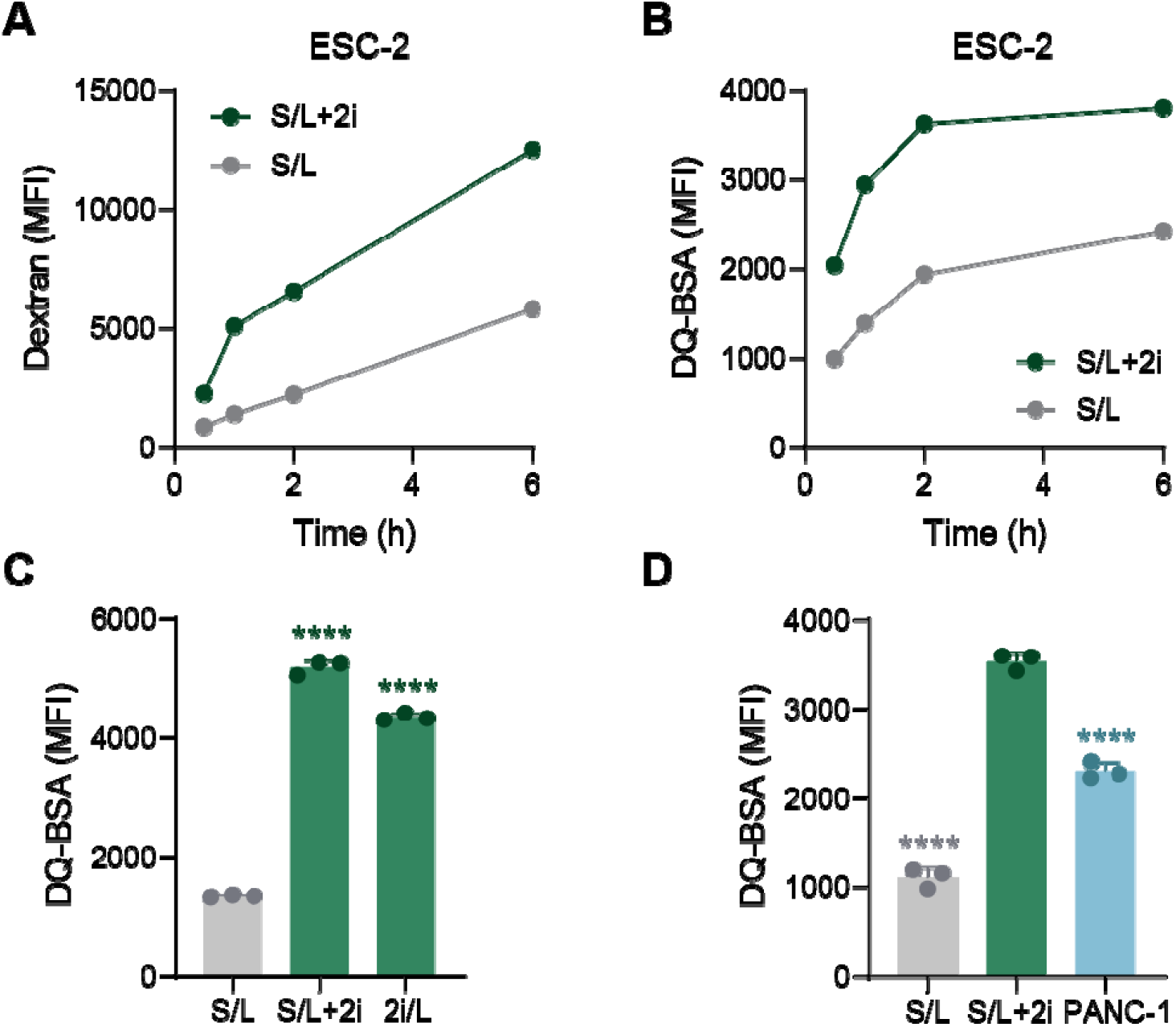
Protein uptake and hydrolysis in ESCs. (**A** and **B**) Dextran uptake (A) and DQ-BSA hydrolysis (B) in S/L and S/L+2i cultured ESCs (ESC-2). (**C**) Flow cytometry quantification of DQ-BSA signal in S/L, S/L+2i and 2i/L-cultured ESCs. (**D**) Flow cytometry quantification of DQ-BSA signal in S/L and S/L+2i cultured ESCs and PANC-1 pancreatic ductal adenocarcinoma cells. Data are single measurements (A and B) or mean ± SD, n=3 independent samples (C and D). Significance was assessed by assessed in comparison to S/L [(C)] or S/L+2i [(D)] cells by one-way ANOVA with Sidak’s multiple comparisons post-test (\*\*\*\**p*<0.0001).

**Supplemental Fig.3.**
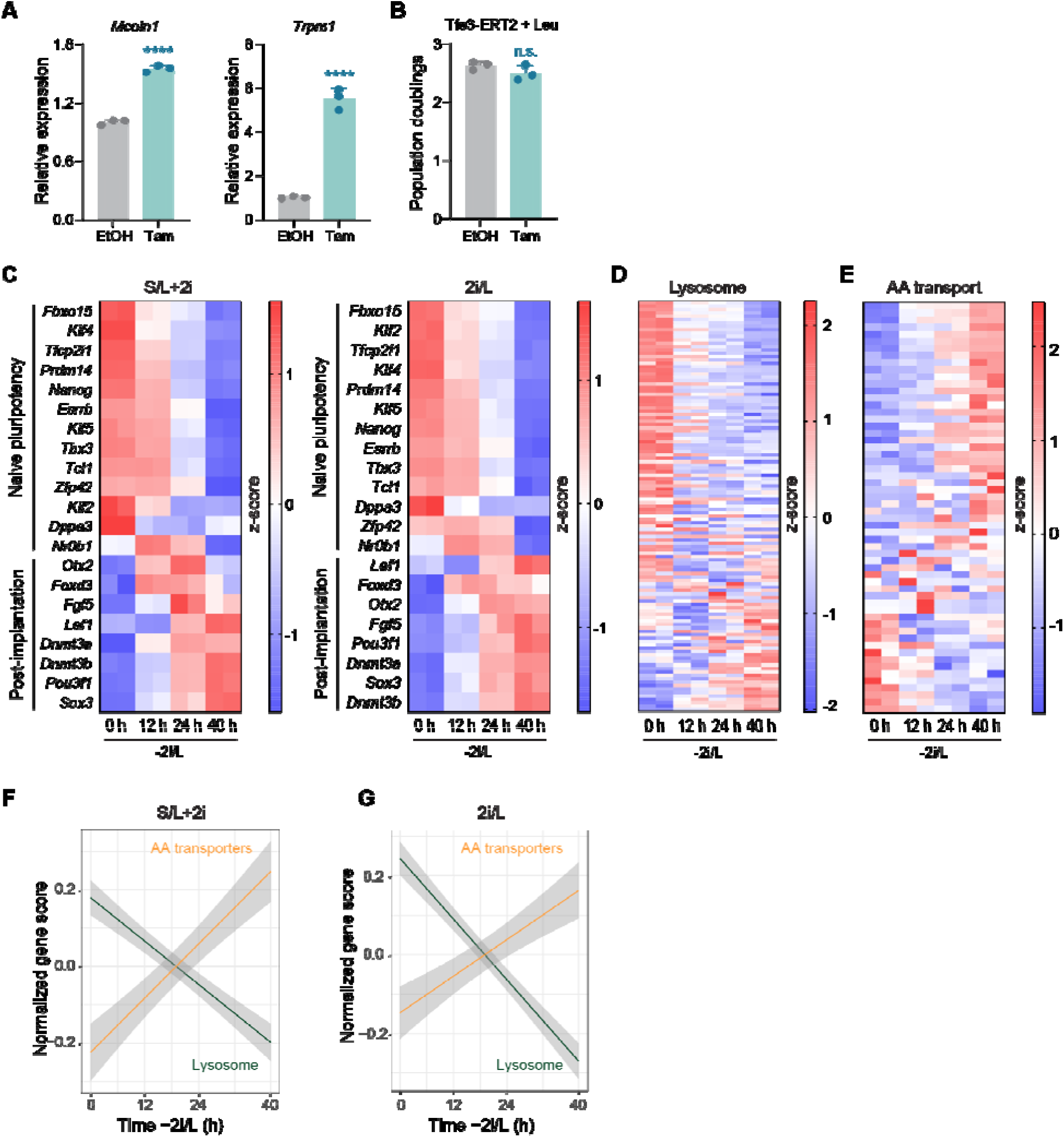
Gene expression changes in ESCs following Tfe3 activation or exit from pluripotency. (**A**) RT-qPCR of Tfe3 target genes (*Mcoln1, Trpm1*) in Tfe3-ERT2-expressing S/L cultured ESCs treated with vehicle (ethanol, EtOH) or tamoxifen (Tam). (**B**) Population doublings of Tfe3-ERT2-expressing S/L cultured ESCs subjected to leucine (Leu) replete conditions for 48 h. (**C-E**) RNA-seq analysis of S/L+2i and 2i/L-cultured ESCs subjected to exit from pluripotency conditions (−2i/L) for 12, 24 and 40 h. Genes encoding naïve pluripotency and early post-implantation are shown. (**D** and **E**) Heat maps showing expression of KEGG lysosome genes (D) or plasma membrane amino acid transporters (E) in S/L+2i cultured ESCs subjected to exit from pluripotency conditions for 12, 24 and 40 h. (**F** and **G**) Regression line through normalized expression of KEGG lysosome genes or plasma membrane amino acid transporters in S/L+2i (F) or 2i/L(G) cultured ESCs subjected to exit from pluripotency conditions for 12, 24 and 40 h. Standard error is shown in grey. Data are mean ± SD, n=3 independent samples (A and B); n=2 independent samples (C-E). Significance was assessed by unpaired two-tailed *t* test [(A-C)] (n.s.=not significant, \*\*\*\**p*<0.0001).

**Supplemental Fig.4.**
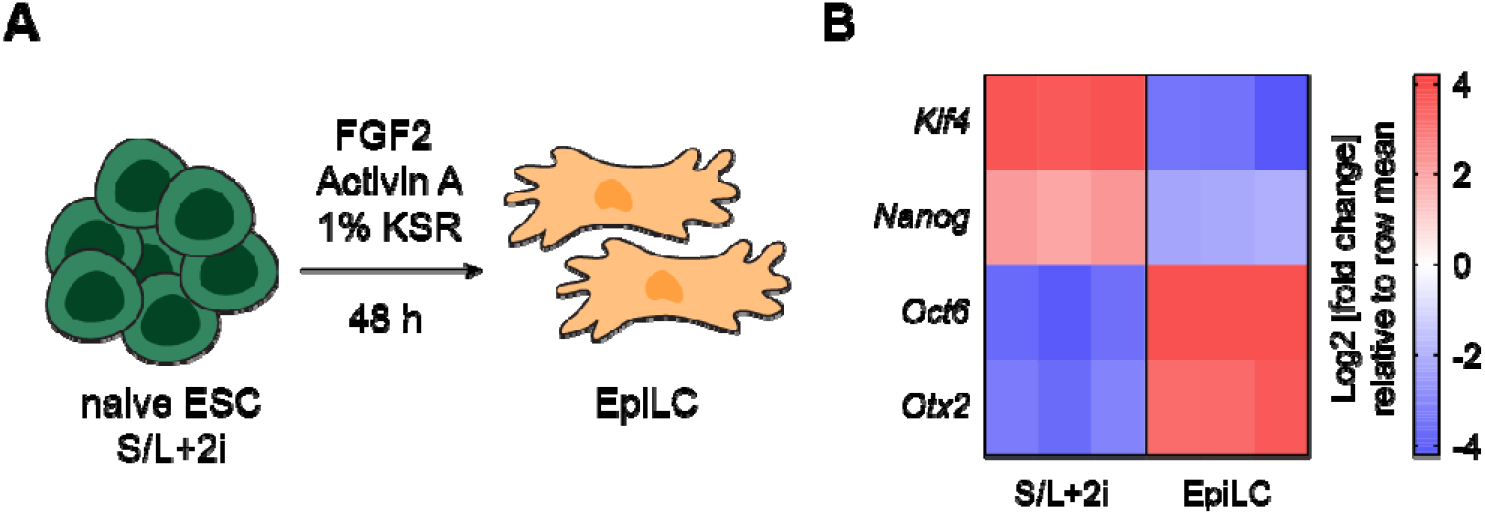
Conversion of naïve ESCs to EpiLCs. (**A**) Schematic of the conversion of S/L+2i cultured ESCs to EpiLCs. (**B**) RT-qPCR of naïve pluripotency-associated (*Klf4, Nanog*) and EpiLC-associated (*Oct6, Otx2*) genes in S/L+-2i cultured ESCs maintained under control or EpiLC-inducing conditions for 48 h. n=3 independent samples.

## Notes

### Competing Interest Statement

The authors have declared no competing interest.

## References

1 Rivera-Perez, J. A. & Hadjantonakis, A. K. The Dynamics of Morphogenesis in the Early Mouse Embryo. Cold Spring Harb Perspect Biol 7, doi:10.1101/cshperspect.a015867 (2014).

2 Alberio, R. Regulation of Cell Fate Decisions in Early Mammalian Embryos. Annu Rev Anim Biosci 8, 377–393, doi:10.1146/annurev-animal-021419-083841 (2020).

3 Brinster, R. L. Effect of glutathione on the development of two-cell mouse embryos in vitro. J Reprod Fertil 17, 521–525, doi:10.1530/jrf.0.0170521 (1968).

4 Whitten, W. K. & Biggers, J. D. Complete development in vitro of the pre-implantation stages of the mouse in a simple chemically defined medium. J Reprod Fertil 17, 399–401, doi:10.1530/jrf.0.0170399 (1968).

5 Cholewa, J. A. & Whitten, W. K. Development of two-cell mouse embryos in the absence of a fixed-nitrogen source. J Reprod Fertil 22, 553–555, doi:10.1530/jrf.0.0220553 (1970).

6 Leese, H. J. Metabolism of the preimplantation embryo: 40 years on. Reproduction 143, 417–427, doi:10.1530/REP-11-0484 (2012).

7 Hosios, A. M. et al. Amino Acids Rather than Glucose Account for the Majority of Cell Mass in Proliferating Mammalian Cells. Dev Cell 36, 540–549, doi:10.1016/j.devcel.2016.02.012 (2016).

8 Palm, W. & Thompson, C. B. Nutrient acquisition strategies of mammalian cells. Nature 546, 234–242, doi:10.1038/nature22379 (2017).

9 Lu, V., Roy, I. J. & Teitell, M. A. Nutrients in the fate of pluripotent stem cells. Cell Metab 33, 2108–2121, doi:10.1016/j.cmet.2021.09.013 (2021).

10 Mathieu, J. & Ruohola-Baker, H. Metabolic remodeling during the loss and acquisition of pluripotency. Development 144, 541–551, doi:10.1242/dev.128389 (2017).

11 Boroviak, T., Loos, R., Bertone, P., Smith, A. & Nichols, J. The ability of inner-cell-mass cells to self-renew as embryonic stem cells is acquired following epiblast specification. Nat Cell Biol 16, 516–528, doi:10.1038/ncb2965 (2014).

12 Ying, Q. L. et al. The ground state of embryonic stem cell self-renewal. Nature 453, 519–523, doi:10.1038/nature06968 (2008).

13 Martello, G. & Smith, A. The nature of embryonic stem cells. Annu Rev Cell Dev Biol 30, 647–675, doi:10.1146/annurev-cellbio-100913-013116 (2014).

14 Hayashi, K., Ohta, H., Kurimoto, K., Aramaki, S. & Saitou, M. Reconstitution of the Mouse Germ Cell Specification Pathway in Culture by Pluripotent Stem Cells. Cell 146, 519–532, doi:10.1016/j.cell.2011.06.052 (2011).

15 Nakamura, T. et al. A developmental coordinate of pluripotency among mice, monkeys and humans. Nature 537, 57–62, doi:10.1038/nature19096 (2016).

16 Kalkan, T. et al. Tracking the embryonic stem cell transition from ground state pluripotency. Development 144, 1221–1234, doi:10.1242/dev.142711 (2017).

17 Arnold, P. K. et al. A non-canonical tricarboxylic acid cycle underlies cellular identity. Nature 603, 477–481, doi:10.1038/s41586-022-04475-w (2022).

18 Carey, B. W., Finley, L. W., Cross, J. R., Allis, C. D. & Thompson, C. B. Intracellular alpha-ketoglutarate maintains the pluripotency of embryonic stem cells. Nature 518, 413–416, doi:10.1038/nature13981 (2015).

19 Vardhana, S. A. et al. Glutamine independence is a selectable feature of pluripotent stem cells. Nat Metab 1, 676–687, doi:10.1038/s42255-019-0082-3 (2019).

20 Bayerl, J. et al. Principles of signaling pathway modulation for enhancing human naive pluripotency induction. Cell Stem Cell 28, 1549–1565 e1512, doi:10.1016/j.stem.2021.04.001 (2021).

21 Xu, Y. et al. Chaperone-mediated autophagy regulates the pluripotency of embryonic stem cells. Science 369, 397–403, doi:10.1126/science.abb4467 (2020).

22 Brune, D., Andrade-Navarro, M. A. & Mier, P. Proteome-wide comparison between the amino acid composition of domains and linkers. BMC Res Notes 11, 117, doi:10.1186/s13104-018-3221-0 (2018).

23 Shyh-Chang, N. et al. Influence of threonine metabolism on S-adenosylmethionine and histone methylation. Science 339, 222–226, doi:10.1126/science.1226603 (2013).

24 Wang, J. et al. Dependence of mouse embryonic stem cells on threonine catabolism. Science 325, 435–439, doi:10.1126/science.1173288 (2009).

25 Commisso, C. et al. Macropinocytosis of protein is an amino acid supply route in Ras-transformed cells. Nature 497, 633–637, doi:10.1038/nature12138 (2013).

26 Lee, S. W. et al. EGFR-Pak Signaling Selectively Regulates Glutamine Deprivation-Induced Macropinocytosis. Dev Cell 50, 381–392 e385, doi:10.1016/j.devcel.2019.05.043 (2019).

27 Davidson, S. M. et al. Direct evidence for cancer-cell-autonomous extracellular protein catabolism in pancreatic tumors. Nature Medicine 23, 235–241, doi:10.1038/nm.4256 (2016).

28 Kamphorst, J. J. et al. Human Pancreatic Cancer Tumors Are Nutrient Poor and Tumor Cells Actively Scavenge Extracellular Protein. Cancer Research 75, 544–553, doi:10.1158/0008-5472.Can-14-2211 (2015).

29 Recouvreux, M. V. & Commisso, C. Macropinocytosis: A Metabolic Adaptation to Nutrient Stress in Cancer. Front Endocrinol (Lausanne) 8, 261, doi:10.3389/fendo.2017.00261 (2017).

30 Reis, R. C., Sorgine, M. H. & Coelho-Sampaio, T. A novel methodology for the investigation of intracellular proteolytic processing in intact cells. Eur J Cell Biol 75, 192–197, doi:10.1016/S0171-9335(98)80061-7 (1998).

31 Kim, S. M. et al. PTEN Deficiency and AMPK Activation Promote Nutrient Scavenging and Anabolism in Prostate Cancer Cells. Cancer Discovery 8, 866–883, doi:10.1158/2159-8290.Cd-17-1215 (2018).

32 West, M. A., Bretscher, M. S. & Watts, C. Distinct endocytotic pathways in epidermal growth factor-stimulated human carcinoma A431 cells. J Cell Biol 109, 2731–2739, doi:10.1083/jcb.109.6.2731 (1989).

33 Montalvo-Ortiz, B. L. et al. Characterization of EHop-016, novel small molecule inhibitor of Rac GTPase. J Biol Chem 287, 13228–13238, doi:10.1074/jbc.M111.334524 (2012).

34 Deacon, S. W. et al. An isoform-selective, small-molecule inhibitor targets the autoregulatory mechanism of p21-activated kinase. Chem Biol 15, 322–331, doi:10.1016/j.chembiol.2008.03.005 (2008).

35 Charpentier, J. C. et al. Macropinocytosis drives T cell growth by sustaining the activation of mTORC1. Nat Commun 11, 180, doi:10.1038/s41467-019-13997-3 (2020).

36 Faddah, D. A. et al. Single-cell analysis reveals that expression of nanog is biallelic and equally variable as that of other pluripotency factors in mouse ESCs. Cell Stem Cell 13, 23–29, doi:10.1016/j.stem.2013.04.019 (2013).

37 Wray, J. et al. Inhibition of glycogen synthase kinase-3 alleviates Tcf3 repression of the pluripotency network and increases embryonic stem cell resistance to differentiation. Nat Cell Biol 13, 838–845, doi:10.1038/ncb2267 (2011).

38 Marks, H. et al. The transcriptional and epigenomic foundations of ground state pluripotency. Cell 149, 590–604, doi:10.1016/j.cell.2012.03.026 (2012).

39 Betschinger, J. et al. Exit from Pluripotency Is Gated by Intracellular Redistribution of the bHLH Transcription Factor Tfe3. Cell 153, 335–347, doi:10.1016/j.cell.2013.03.012 (2013).

40 Nowotschin, S. et al. The emergent landscape of the mouse gut endoderm at single-cell resolution. Nature 569, 361–367, doi:10.1038/s41586-019-1127-1 (2019).

41 Hayashi, K. & Saitou, M. Stepwise differentiation from naive state pluripotent stem cells to functional primordial germ cells through an epiblast-like state. Methods Mol Biol 1074, 175–183, doi:10.1007/978-1-62703-628-3_13 (2013).

42 Shao, X. et al. Placental trophoblast syncytialization potentiates macropinocytosis via mTOR signaling to adapt to reduced amino acid supply. Proceedings of the National Academy of Sciences 118, e2017092118, doi:10.1073/pnas.2017092118 (2021).

43 Atlasi, Y. et al. Epigenetic modulation of a hardwired 3D chromatin landscape in two naive states of pluripotency. Nature Cell Biology 21, 568–578, doi:10.1038/s41556-019-0310-9 (2019).

44 Hamilton, T. G., Klinghoffer, R. A., Corrin, P. D. & Soriano, P. Evolutionary divergence of platelet-derived growth factor alpha receptor signaling mechanisms. Mol Cell Biol 23, 4013–4025, doi:10.1128/MCB.23.11.4013-4025.2003 (2003).

45 Plusa, B., Piliszek, A., Frankenberg, S., Artus, J. & Hadjantonakis, A. K. Distinct sequential cell behaviours direct primitive endoderm formation in the mouse blastocyst. Development 135, 3081–3091, doi:10.1242/dev.021519 (2008).

46 Brison, D. R. et al. Identification of viable embryos in IVF by non-invasive measurement of amino acid turnover. Hum Reprod 19, 2319–2324, doi:10.1093/humrep/deh409 (2004).

47 Houghton, F. D. et al. Non-invasive amino acid turnover predicts human embryo developmental capacity. Hum Reprod 17, 999–1005, doi:10.1093/humrep/17.4.999 (2002).

48 Leese, H. J., Brison, D. R. & Sturmey, R. G. The Quiet Embryo Hypothesis: 20 years on. Front Physiol 13, 899485, doi:10.3389/fphys.2022.899485 (2022).

49 Hoijman, E. et al. Cooperative epithelial phagocytosis enables error correction in the early embryo. Nature 590, 618–623, doi:10.1038/s41586-021-03200-3 (2021).

50 Whitten, W. K. in Schering Symposium on Intrinsic and Extrinsic Factors in Early Mammalian Development, Venice, April 20 to 23, 1970 (ed Gerhard Raspé) 129–141 (Pergamon, 1971).

51 Sonder, S. L. et al. Restructuring of the plasma membrane upon damage by LC3-associated macropinocytosis. Sci Adv 7, doi:10.1126/sciadv.abg1969 (2021).

52 Kostopoulou, N. et al. Embryonic stem cells are devoid of macropinocytosis, a trafficking pathway for activin A in differentiated cells. J Cell Sci 134, doi:10.1242/jcs.246892 (2021).

53 von Zastrow, M. & Sorkin, A. Mechanisms for Regulating and Organizing Receptor Signaling by Endocytosis. Annu Rev Biochem 90, 709–737, doi:10.1146/annurev-biochem-081820-092427 (2021).

54 Villegas, F. et al. Lysosomal Signaling Licenses Embryonic Stem Cell Differentiation via Inactivation of Tfe3. Cell Stem Cell 24, 257–270 e258, doi:10.1016/j.stem.2018.11.021 (2019).

55 Puccini, J., Badgley, M. A. & Bar-Sagi, D. Exploiting cancer’s drinking problem: regulation and therapeutic potential of macropinocytosis. Trends Cancer 8, 54–64, doi:10.1016/j.trecan.2021.09.004 (2022).

56 Vander Heiden, M. G. & DeBerardinis, R. J. Understanding the Intersections between Metabolism and Cancer Biology. Cell 168, 657–669, doi:10.1016/j.cell.2016.12.039 (2017).

57 Lu, V. et al. Glutamine-dependent signaling controls pluripotent stem cell fate. Dev Cell 57, 610–623 e618, doi:10.1016/j.devcel.2022.02.003 (2022).

58 Young, N. P. et al. AMPK governs lineage specification through Tfeb-dependent regulation of lysosomes. Genes Dev 30, 535–552, doi:10.1101/gad.274142.115 (2016).

59 Morgani, S. M. & Hadjantonakis, A. K. Spatially Organized Differentiation of Mouse Pluripotent Stem Cells on Micropatterned Surfaces. Methods Mol Biol 2214, 41–58, doi:10.1007/978-1-0716-0958-3_4 (2021).

60 Chen, S., Zhou, Y., Chen, Y. & Gu, J. fastp: an ultra-fast all-in-one FASTQ preprocessor. Bioinformatics 34, i884–i890, doi:10.1093/bioinformatics/bty560 (2018).

61 Liao, Y., Smyth, G. K. & Shi, W. featureCounts: an efficient general purpose program for assigning sequence reads to genomic features. Bioinformatics 30, 923–930, doi:10.1093/bioinformatics/btt656 (2014).

62 Love, M. I., Huber, W. & Anders, S. Moderated estimation of fold change and dispersion for RNA-seq data with DESeq2. Genome Biol 15, 550, doi:10.1186/s13059-014-0550-8 (2014).

63 Wickham, H. in Use R!, 1 online resource (XVI, 260 pages 232 illustrations, 140 illustrations in color (Springer International Publishing : Imprint: Springer,, Cham, 2016).

64 Subramanian, A. et al. Gene set enrichment analysis: a knowledge-based approach for interpreting genome-wide expression profiles. Proc Natl Acad Sci U S A 102, 15545–15550, doi:10.1073/pnas.0506580102 (2005).

65 Wu, T. et al. clusterProfiler 4.0: A universal enrichment tool for interpreting omics data. Innovation (Camb) 2, 100141, doi:10.1016/j.xinn.2021.100141 (2021).

66 van Dijk, D. et al. Recovering Gene Interactions from Single-Cell Data Using Data Diffusion. Cell 174, 716–729 e727, doi:10.1016/j.cell.2018.05.061 (2018).

67 Hao, Y. et al. Integrated analysis of multimodal single-cell data. Cell 184, 3573–3587 e3529, doi:10.1016/j.cell.2021.04.048 (2021).

68 Commisso, C., Flinn, R. J. & Bar-Sagi, D. Determining the macropinocytic index of cells through a quantitative image-based assay. Nature Protocols 9, 182–192, doi:10.1038/nprot.2014.004 (2014).

69 Solter, D. & Knowles, B. B. Immunosurgery of mouse blastocyst. Proc Natl Acad Sci U S A 72, 5099–5102, doi:10.1073/pnas.72.12.5099 (1975).

